# Tissue resident memory CD4^+^ T cells are sustained by site-specific levels of self-renewal and continuous replacement

**DOI:** 10.1101/2024.09.26.615039

**Authors:** Jodie Chandler, M. Elise Bullock, Arpit C. Swain, Cayman Williams, Christiaan H. van Dorp, Benedict Seddon, Andrew J. Yates

## Abstract

Tissue-resident memory T cells (T_RM_) protect from repeat infections within organs and barrier sites. The breadth and duration of such protection is defined at minimum by three quantities; the rate at which new T_RM_ are generated from precursors, their rate of self-renewal, and their rate of loss through death, egress, or differentiation. Quantifying these processes individually is challenging. Here we combine genetic fate mapping tools and mathematical models to untangle these basic homeostatic properties of CD4^+^ T_RM_ in the skin and gut lamina propria (LP) of healthy adult mice. We show that CD69^+^CD4^+^ T_RM_ in skin reside for ∼24 days and self-renew more slowly, such that clones halve in size approximately every 5 weeks; and approximately 2% of cells are replaced daily from precursors. CD69^+^CD4^+^ T_RM_ in LP have shorter residencies (∼14 days) and are maintained largely by immigration (4-6% per day). We also find evidence that the continuous replacement of CD69^+^CD4^+^ T_RM_ at both sites derives from circulating effector-memory CD4^+^ T cells, in skin possibly via a local CD69^−^ intermediate. Our approach maps the ontogeny of CD4^+^ T_RM_ in skin and LP and exposes their dynamic and distinct behaviours, with continuous seeding and erosion potentially impacting the duration of immunity at these sites.

## Introduction

Resident memory T cells (T_RM_) provide immune surveillance and protection in tissues throughout the body (Szabo *et al*., 2019), but the mechanisms by which they are maintained are not well understood. Conventional CD4^+^ and CD8^+^ T_RM_ in mice and humans are not intrinsically long-lived, but appear to self-renew slowly as assessed by readouts of cell division such Ki67 expression or BrdU incorporation, at levels that vary across tissues (Gebhardt *et al*., 2009, Watanabe *et al*., 2015, Park *et al*., 2018, Strobl *et al*., 2020, Divito *et al*., 2020, Christo *et al*., 2021). After infection or immune challenge, the numbers of elicited T_RM_ may also be sustained by influx from precursor populations, although the extent to which this occurs is unclear, and is likely also cell subset- and tissue-dependent. For example, in the lung there is evidence both for (Zammit *et al*., 2006, Ely *et al*., 2006, Slütter *et al*., 2017, Van Braeckel-Budimir *et al*., 2018, Takamura and Kohlmeier, 2019) and against (Takamura *et al*., 2016, van Dorp *et al*., 2024) ongoing recruitment of new T_RM_ following respiratory virus infections. Within skin, T_RM_ may be renewed or supplemented slowly from precursors in the setting of graft-versus-host disease (Divito *et al*., 2020), and from circulating central memory T_CM_ or effector memory T_EM_ following infection or sensitisation (Gaide *et al*., 2015, Matos *et al*., 2022). In the small intestine, however, resident CD4^+^ and CD8^+^ T_RM_ appear to persist for months to years with slow self-renewal without appreciable influx (Bartolomé-Casado *et al*., 2019, 2021).

The dynamics of production and loss of T_RM_ in the steady state are even less well understood, and measuring these processes is important for several reasons. The balance of loss and self-renewal defines the persistence of clonal populations and hence the duration of protective immunity. Further, while self-renewal can at best preserve clonal diversity within a tissue site, any supplementation or replacement by immigrant T_RM_ will perturb the local TCR repertoire. In particular, any significant influx into T_RM_ niches in the absence of overt infection may be a competitive force, potentially reducing the persistence T_RM_ previously established in response to infection or challenge.

The kinetics of circulating memory T cells have been quantified extensively in both mice and humans, using dye dilution assays (Choo *et al*., 2010), deuterium labelling (Westera *et al*., 2013, 2015, del Amo *et al*., 2018, Baliu-Piqué *et al*., 2018, 2019, van den Berg *et al*., 2021), and BrdU labelling, either alone (Younes *et al*., 2011, Ganusov and De Boer, 2013) or in combination with fate reporters (Gossel *et al*., 2017, Hogan *et al*., 2019, Bullock *et al*., 2024) or T cell receptor excision circles (den Braber *et al*., 2012). Using mathematical models to interpret these data, these studies identified rates of production, cellular lifespans, and signatures of heterogeneity in turnover. modelling has also established evidence for continuous replenishment of circulating memory CD4^+^ T cells from precursors throughout life in specific-pathogen-free mice, driven by a combination of environmental, commensal and self antigens (Gossel *et al*., 2017, Hogan *et al*., 2019, Bullock *et al*., 2024). Quantification of T_RM_ dynamics has to date been restricted largely to measuring the net persistence of CD8^+^ T_RM_ following infection in mice, in a variety of tissues (Morris *et al*., 2019, Wijeyesinghe *et al*., 2021). Far less is known regarding CD4^+^ T_RM_, which typically outnumber their CD8^+^ counterparts (Szabo *et al*., 2019), and there have been very few attempts to dissect the kinetics of either subset (van Dorp *et al*., 2024).

In general, measuring these basic parameters in isolation is challenging, partly due to their sensitivity to assumptions made in the models (De Boer and Perelson, 2013), but also because division-linked labelling alone may not distinguish *in situ* cell division and the supplementation of a population from labelled precursors. A more powerful approach is to triangulate information from different readouts of cell fate simultaneously (Bains *et al*., 2009, den Braber *et al*., 2012, del Amo *et al*., 2018, De Boer and Yates, 2023, Bullock *et al*., 2024).

With these challenges in mind, here we integrated data from two independent inducible fate reporter systems to study CD4^+^ T_RM_ homeostasis in mice. Each system allows one to track the fates of defined populations of cells and their descendants. One labels all CD4^+^ T cell subsets at any given moment, which effectively provides an age ‘timestamp’. The other labels cells that are dividing during a defined time window. In combination, these systems allowed us to establish a quantitative model of the basal homeostatic properties of CD4^+^ T_RM_ within the skin and the lamina propria of the small intestine in healthy mice. In particular, we could unpick the contributions of self renewal and *de novo* cell production that underpin their maintenance, and explore their relationships to circulating T cell subsets.

## Results

### Combining cell fate reporters and models to measure T_RM_ replacement, loss, and self-renewal

To study the homeostatic dynamics of tissue resident CD4^+^ memory T cells in healthy mice, we used in concert two genetic fate mapping tools in which cohorts of peripheral T cells and their offspring can be induced to express permanent fluorescent markers (Fig. 1A). These reporter strains were previously used separately to study the turnover of naive and circulating memory T and B cells (Verheijen *et al*., 2020, Lukas *et al*., 2023, Bullock *et al*., 2024). In the Ki67^mCherry-CreERT^ Rosa26^RcagYFP^ system, henceforth Ki67-DIVN, fluorescent reporters are linked to the expression of Ki67, a nuclear protein that is expressed during cell division and for 3 to 4 days afterwards (Gossel *et al*., 2017, Miller *et al*., 2018). Specifically, these mice express both a Ki67-mCherry fusion protein and inducible CreERT from the *Mki67* locus, together with a *Rosa26*^*RcagYFP*^ Cre reporter construct. Treatment of mice with tamoxifen therefore induces YFP in cells expressing high levels of Ki67, and YFP is then stably expressed by these cells and their offspring. Expression of Ki67-fused mCherry gives a constitutive live readout of Ki67 expression, independent of tamoxifen treatment and YFP expression. In the second fate reporter, CD4^CreERT^ Rosa26^RmTom^ mice (Cd4-FR), the Cre reporter is constructed such that cells expressing CD4 during tamoxifen treatment permanently and heritably express the fluorescent reporter mTomato (mTom).

**Figure 1.**
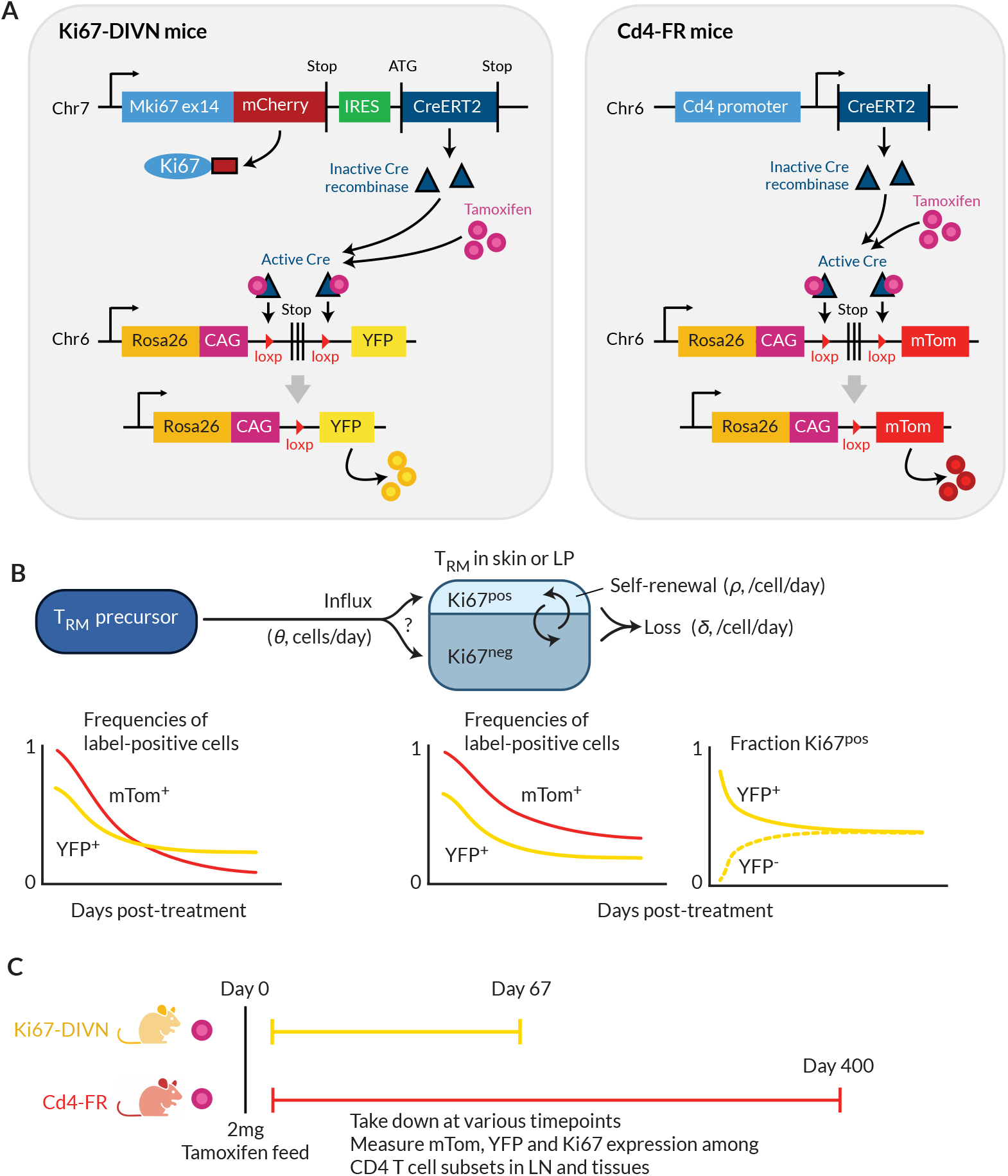
Overview of the fate-reporter approach to quantifying the homeostasis of skin- and LP-resident CD4^+^ T cells. **A** Ki67 and CD4 reporter constructs. **B** Schematic of a simple mathematical model of TRM homeostasis at steady state. New cells enter a TRM subset in skin or LP (the ‘target’ population) from a precursor population at rate *θ* (cells per unit time). YFP and mTom expression among immigrant cells is assumed to be that of the precursor population in bulk. We considered three possibilities for the Ki67 expression levels among new immigrants; a ‘quiescent’ mode (all Ki67^low^), ‘neutral’ recruitment (Ki67^high^ frequency identical to that of the precursor), or ‘division-linked’ (all recruited cells Ki67^high^). The target cells are assumed to be a kinetically homogeneous population that self-renews at average rate *ρ*, such that their mean interdivision time is 1*/ρ*. Cells are also lost through death, differentiation or tissue egress at total rate δ, implying a mean residence time of 1*/*δ. The combination of these processes determines the timecourses of frequencies of YFP^+^ and mTom^+^ cells with the target population. **C** Experimental design.

In a closed population of cells at steady state, self-renewal must be balanced by loss and so, following tamoxifen treatment, the frequencies of cells expressing YFP or mTom within any such population would remain constant. Therefore, any decline in the frequency of either reporter among T_RM_ after treatment must derive from the influx of label-negative cells from an upstream (precursor) population. In the Ki67-DIVN mice, these will be descendants of cells that were not dividing at the time of tamoxifen treatment; in the Cd4-FR mice, labelled CD4^+^ cells will slowly be replaced by the descendants of those generated in the thymus after treatment. The shape of this decline will be determined by the combination of the net loss rate of T_RM_ from the tissue (through death, egress, or differentiation, offset by any self-renewal), and the label content of immigrant T_RM_ (Fig. 1B). To refer to the persistence of individual T_RM_ cells we will use the term ‘residence time’ rather than lifespan, to reflect the multiple potential mechanisms of loss from tissues.

To quantify these processes, we treated cohorts of both reporter mice, aged between 4 and 15 weeks, with a single 2mg pulse of tamoxifen (Fig. 1C). Over 9 week (Ki67 reporter) and 57 week (CD4 reporter) chase periods we measured the frequencies of labelled cells among antigen-experienced CD4^+^ T cell subsets isolated from skin and the lamina propria of the small intestine (henceforth LP), and within circulating naive and memory T cell subsets derived from lymph nodes (Fig. S1A). By combining these frequencies with measures of Ki67 expression, and describing the resulting set of time series with simple mathematical models, we aimed to estimate the basic parameters underlying T_RM_ kinetics.

### CD4+ T_RM_ in skin and lamina propria are continuously replaced from precursors

We considered two populations within both skin and LP, identified as tissue-localised by virtue of their protection from short-term *in vivo* labelling (Methods; Fig. S1B). One was effector-memory (EM) phenotype (CD4^+^CD44^hi^ CD62L^lo^) T cells in bulk, which we studied in order to gain the broadest possible picture of memory T cell dynamics at these sites. We also considered the subset of these cells that expressed CD69, a canonical and consistent marker of CD4^+^ T cell residency across multiple tissues (Szabo *et al*., 2019). We saw no significant changes with mouse age in the numbers of either population within skin (Fig. 2A, *p >*0.67) or LP (Fig. 2B, *p >* 0.39). There were also no significant changes in any of these quantities with time since tamoxifen treatment (*p >* 0.24). We therefore assumed that the skin- and LP-localised T cell subsets we considered were at, or close to, homeostatic equilibrium during the chase period. For brevity, we refer to tissue-localised CD4^+^CD44^hi^CD62L^lo^ in bulk as EM, and their CD69^+^ subset as T_RM_.

**Figure 2.**
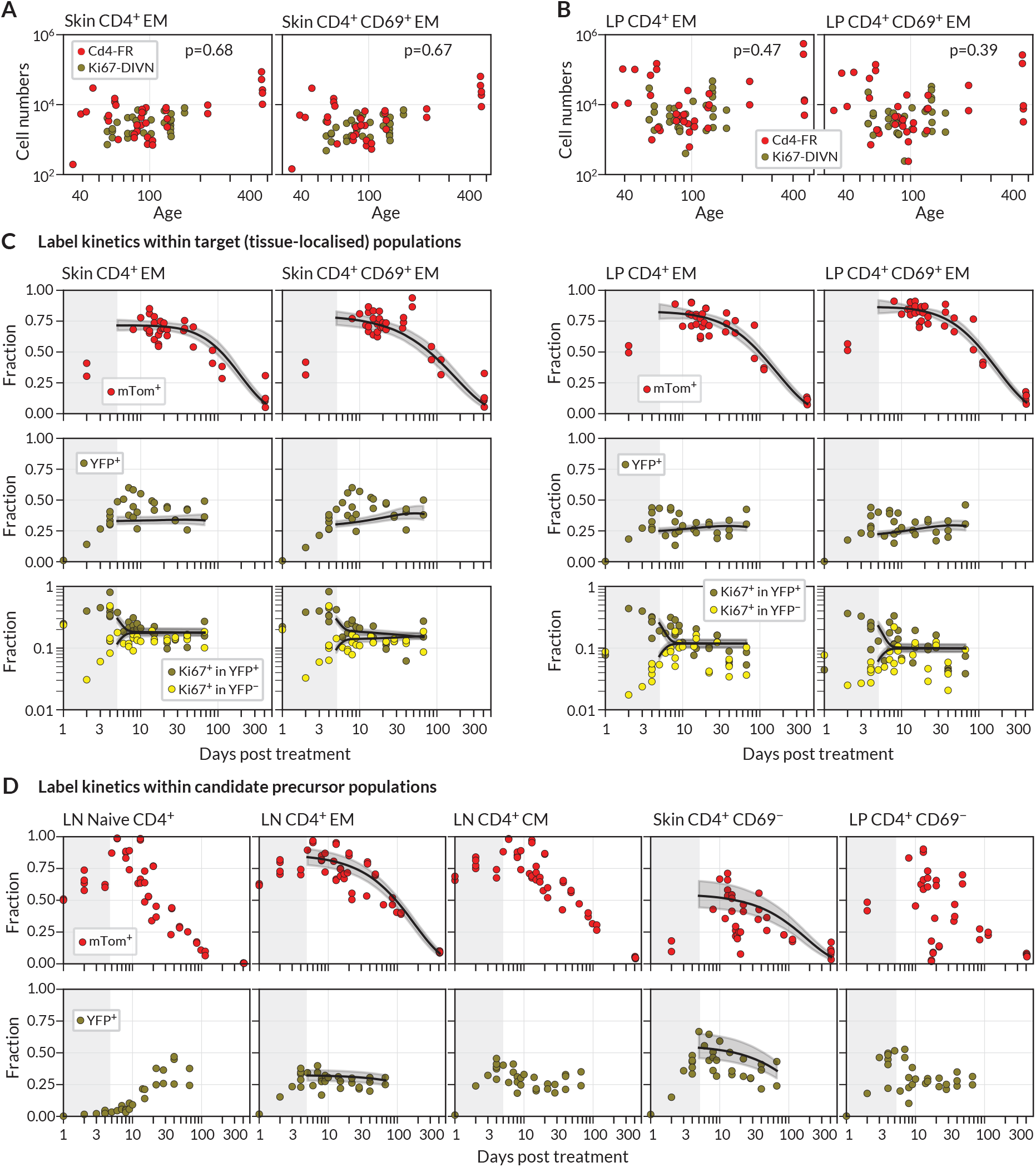
Modelling fate reporter dynamics among CD4^+^ T cell subsets in skin and lamina propria. **A**,**B** Numbers of CD4^+^ effector-memory (EM) phenotype and CD4^+^CD69^+^ T cells recovered from (A) ear skin, and (B) lamina propria in the small intestine, against mouse age. P-values are derived from Spearman rank correlation on data from the Cd4-FR and Ki67-DIVN strains combined. **C** Observed (points), best-fit model trajectories (black lines), and 95% credible intervals (grey envelopes) of the frequencies of mTom^+^ and YFP^+^ CD4^+^ T cell subsets within skin and LP, and the proportions of YFP^+^ and YFP^−^ cells that expressed Ki67, with time since tamoxifen treatment. Shaded rectangles indicate the 5-day period over which reporter expression was being induced; data from these periods were not used in fitting. **D** Frequencies of mTom^+^ and YFP^+^ cells within lymph node CD4^+^ T cell subsets, and within CD4^+^CD44^+^CD69^−^ T cells in skin and LP. Overlaid are the empirical descriptions of these trajectories from the best fitting model in which that population was identified as a precursor (Text S1); grey envelopes are 95% credible intervals. (The naive and CM in lymph nodes, and CD69^−^ cells in LP, were never identified as favoured precursors).

During the first few days after tamoxifen treatment, YFP and mTom expression increased continuously within the skin and LP subsets (Fig. 2C), as well as among CD4^+^ naive (CD44^lo^ CD62L^hi^), central memory (T_CM_, CD44^hi^ CD62L^hi^) and effector memory (T_EM_, CD44^hi^ CD62L^lo^) T cells recovered from lymph nodes (Fig. 2D). These initial increases were driven in part by the intracellular dynamics of the induction of the fluorescent reporters. We therefore began our analyses at day 5 post-treatment, by which time induction was considered complete and the subsequent trajectories of label frequencies reflected only the dynamic processes of cell production and loss. A key observation was that mTom expression within the skin and LP subsets then declined slowly (roughly 7-to 8-fold over the course of a year, Fig. 2C), indicating immediately that these populations were being continuously replaced from precursors. Early in the chase period YFP^+^ T_RM_ expressed Ki67 at higher levels than YFP^−^ cells, as expected, but Ki67 expression in the two populations converged at later times. We return to the interpretation of these kinetics below.

We then investigated the extent to which the simple model illustrated in Fig. 1B could explain these trajectories. Given the observation of continued recruitment, any time-variation in the label content of T_RM_ precursors might leave an imprint on the label kinetics of skin or LP T_RM_ themselves, and thereby help us to identify their developmental pathways. We reasoned that plausible T_RM_ precursors might be LN-derived CD4^+^ naive, T_CM_ or T_EM_; we also considered the possibilities that CD69^−^ cells within skin and LP are the direct precursors of the local CD69^+^ populations. Therefore, we used empirical functions to describe the timecourses of the frequencies of YFP^+^ and mTom^+^ cells within these populations (Fig. 2D), and used these to represent the label composition of cells entering the tissue subsets.

For each tissue subset (‘target’) and precursor pair, we fitted the model simultaneously to six timecourses; the frequencies of (i) YFP expression and (ii) mTom expression among target cells, the proportions of Ki67^high^ cells among (iii) YFP^+^ and (iv) YFP^−^ target cells, and the (v) YFP and (vi) mTom expression kinetics within the precursor (Methods, and Supporting Information Text S1). For each precursor/target pair we considered three modes of influx – one in which new immigrant T_RM_ are Ki67^low^ (‘quiescent’ recruitment); another in which their Ki67 expression directly reflects that of the precursor (‘neutral’ recruitment), and a third in which immigrants have recently divided (Ki67^high^), perhaps through an antigen-driven process (‘division-linked’ recruitment).

### Skin and LP CD4^+^ T_RM_ have similar residence times but exhibit distinct contributions of replacement and self-renewal

For each combination of target population, potential precursor, and potential mode of recruitment, we were able to estimate rates of influx, mean residence times, and mean interdivision times for the target population (Fig. 3 and Table S1; prior and posterior distributions of the parameters of the best fitting models are shown in Fig. S2). The mean residence times of both EM and T_RM_ within skin and LP were ∼3 weeks and 2 weeks respectively. The means of production of new cells differed at the two sites, however. In skin, around 2% of both populations were replaced daily by influx, comparable to the rates of constitutive replacement of circulating memory CD4^+^ T cell subsets (Gossel *et al*., 2017, Hogan *et al*., 2019, Bullock *et al*., 2024), and EM and T_RM_ self-renewed every 6 and 7 weeks respectively, In contrast, within LP these subsets divided less often (every 7-9 weeks) and relied on higher levels of recruitment (4-6% per day) for their maintenance.

**Figure 3.**
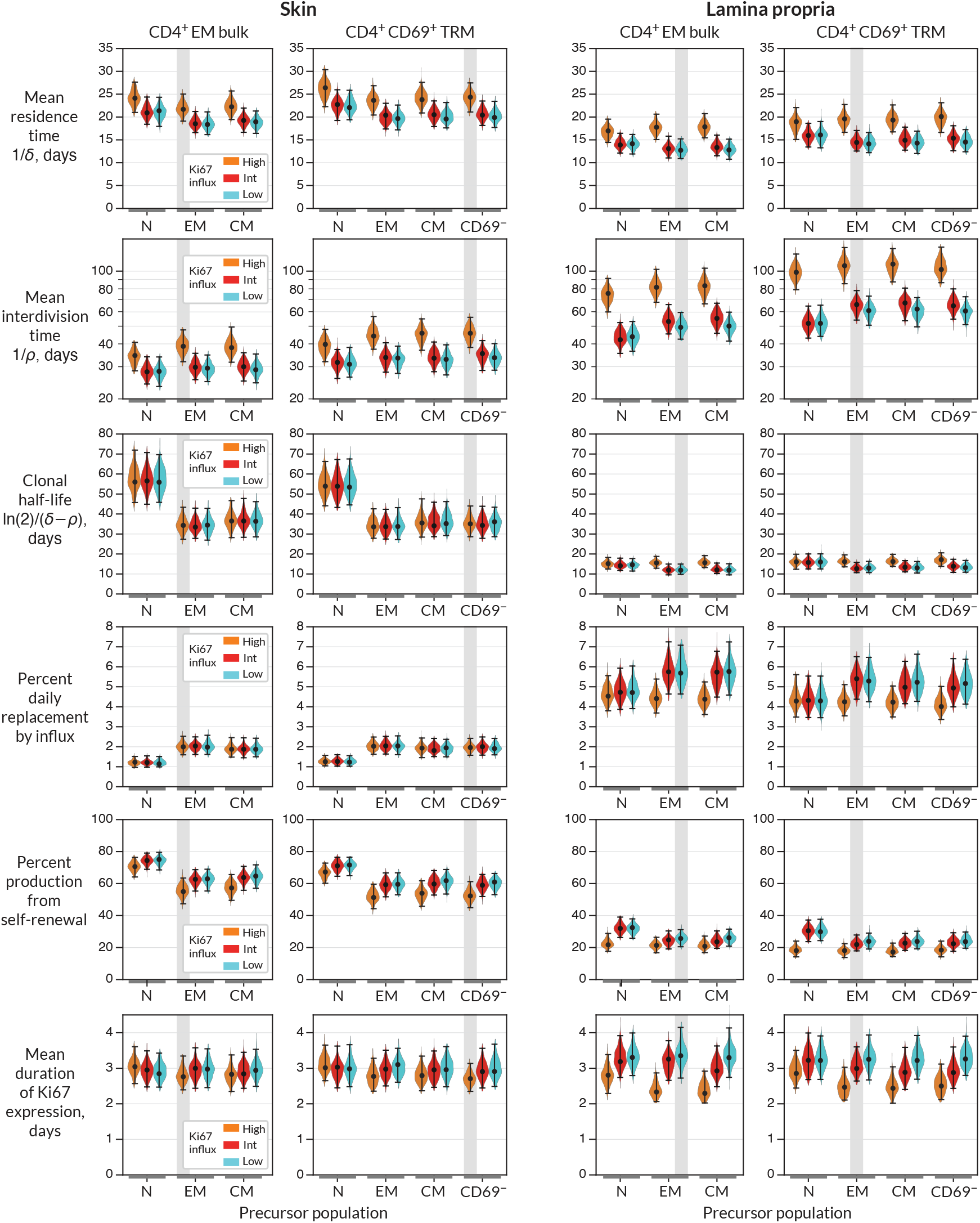
Parameters governing the homeostasis of antigen-experienced CD4^+^ T cells localised within skin and lamina propria (LP) in adult mice. Violin plots indicate the posterior distributions of parameters. Black points and bars; best (maximum *a posteriori*) estimates and 95% credible intervals. For each population (target) in skin or LP, potential precursors were; from lymph nodes, CD4^+^ naive, central memory and effector memory T cells (N, EM and CM); and for CD4^+^ TRM, CD69^+^ TRM within the same tissue. For each precursor/target pair, we considered three potential levels of Ki67 expression on immigrant cells (L-R; orange, red, and blue violin plots). ‘High’, the division-linked model (new cells enter as Ki67^high^); ‘Int’, neutral model (Ki67 expression among new cells mirrors that of the precursor); and ‘Low’, the quiescent model (new cells enter as Ki67^low^). Shaded regions highlight the parameter estimates derived from the favoured model (precursor and Ki67 levels on immigration) for each target (Fig. 4A).

From these basic quantities we could derive several other important measures of T_RM_ behaviour. First, the balance of the rates of loss (δ) and self-renewal (*ρ*) defines the persistence of a cohort of T cells, which is distinct from the lifepan of its constituent cells (del Amo *et al*., 2018, De Boer and Yates, 2023). Specifically, the quantity ln(2)*/*(δ − *ρ*) is the average time taken for a cohort and their descendents to halve in number. While we studied polyclonal populations here, this quantity applies equally well to measuring the persistence of a TCR clonotype, so we refer to it as a clonal half life (Bullock *et al*., 2024). The substantial rates of self-renewal in skin led to clonal half lives of just over a month. The lower levels of self-renewal and higher levels of replacement in LP resulted in shorter clonal half lives of ∼2 weeks.

Importantly, our estimates of these quantities depended to varying degrees on the choice of precursor and mode of recruitment (Fig. 3). For example, intuitively, given the observed level of Ki67 within each target population, the greater the levels of Ki67 within newly recruited cells, the less must derive from self-renewal within the tissue; hence, if one assumes that recruitment is division-linked, estimated division rates are reduced. Similarly, as discussed above, the label content of the precursor influences the net loss rate of label in the target, which was most clearly reflected in the loss of mTom^+^ cells over the longer chase period (Fig. 2C and D). For example, mTom was lost most rapidly within naive CD4^+^ T cells (Fig. 2D), due to export of label-negative cells from the thymus. Models in which naive T cells were the direct precursor of T_RM_ therefore predicted greater clonal persistence within tissues.

We saw very little decline in YFP expression levels during the 2 month chase period (Fig. 2C), due in part to the sustained levels of YFP within the putative precursor populations (Fig. 2D), which ‘topped up’ YFP-expressing T_RM_. As a result, YFP kinetics within the tissues was not strongly informative regarding rates of replacement. However, the rate of convergence of Ki67 within YFP^+^ and YFP^−^ cells (Fig. 2C) put clear constraints on the duration of Ki67 expression, which at approximately 3 days (Fig. 3) was consistent with previous estimates. This quantity in turn was informative for estimating rates of self-renewal. Further, the observed convergence of Ki67 within these two subsets is consistent with the basic model assumption of homogeneity in the rates of division and loss within skin and LP.

### CD4^+^ CD69^+^ T_RM_ within LP likely derive predominantly from circulating T_EM_ in lymph nodes, while those in skin may derive from a CD69^−^ intermediate

To more precisely quantify the kinetics of each target population, we assessed the relative support for each combination of precursor and mode of recruitment (Fig. 4A). Each of these weights summarises the magnitude and uncertainty of a model’s out-of-sample prediction error of the label kinetics within the target population (Methods).

**Figure 4.**
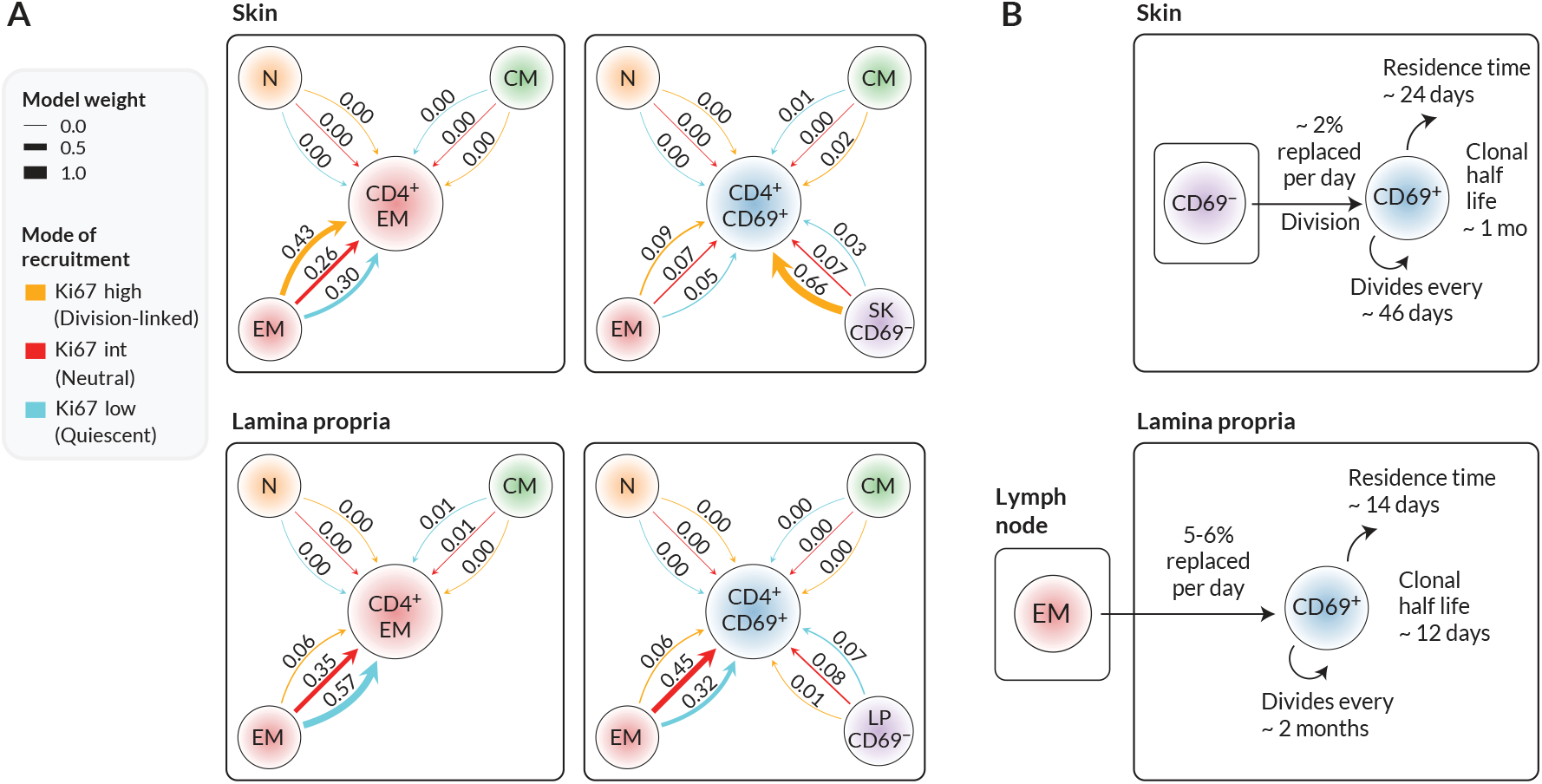
Candidate ontogenic pathways of CD4^+^ CD69^+^ TRM in skin and lamina propria. **A** For each target population with skin and LP (CD4^+^ EM bulk, and CD4^+^CD69^+^ T cells), we calculated the relative support for alternative pathways and modes of recruitment, using LOO-IC weights (Methods). For potential precursors, N, CM, EM refer to naive, TCM and TEM in lymph nodes; for CD69^+^ targets we also considered CD69^−^ cells within the same tissue as a possible precursor. As described in the text, for each precursor/target pair, there were three potential submodels relating to the possible levels of Ki67 on newly recruited cells. Model weights sum to 1 for each target population, with the width of each arrow reflecting a model’s degree of support. **B** Schematics of the most strongly supported pathways of development of CD4^+^CD69^+^ TRM in skin and LP, with approximate values of key kinetic parameters (Fig. 3 and Table S1).

We found that the data were quite strongly informative regarding the immediate ancestors of tissue subsets. From the candidate set of models, the weighting strongly favoured lymph-node derived EM as the closest precursor to CD4^+^ EM within both skin and LP. However, within skin, local CD69^−^ cells were the favoured precursor to CD69^+^ T_RM_. The evidence was generally more equivocal regarding the mode in which cells are recruited into skin and LP, although for skin we saw substantial evidence (66% of model support) for a division-linked transition from CD69^−^ to CD69^+^ cells. Fig. 4B summarises the developmental trajectories and kinetics of T_RM_ in skin and LP that were supported most strongly by our analyses. Parameter estimates and credible intervals for these models are highlighted with vertical shaded regions in Fig. 3 and are detailed in Supporting Information, Table S1. Visual differences between models are shown in Fig. S3, where for each target population we overlay the fits from top-ranked, second-ranked, and lowest-ranked models.

### Validation of residence times through use of Ki67 expression directly

As a consistency check, when a population is at or close to steady state, bounds on the mean residence time of cells can be estimated using only the measured frequency of Ki67 within the target population, and the daily rate of replacement (Supporting Information, Text S2). In skin, both EM and T_RM_ are replaced at the rate of 2% per day, and have Ki67 expression frequencies of around 0.15 (Fig. 2C). The approximation then yields residence times in the range 22-28 days, depending on whether immigrant T_RM_ are Ki67^low^ or Ki67^high^ respectively. In LP, with 5.5% daily replacement and Ki67 frequencies of around 0.07, we estimate residence times of 14-25 days. Both estimates are in good agreement with those from the model fitting. Further validation of these results, and dissection of the kinetics, might be achieved by manipulating cell trafficking, although this would potentially impact multiple processes at once. For example, treating mice with the sphingosine 1-phosphate receptor agonist FTY720 would block tissue ingress and egress. This would leave self-renewal as the only means of T_RM_ production, and would also remove the component of the cell loss rate that is due to cells leaving the tissue. In principle one could then gain estimates of the intrinsic lifespan of T_RM_, rather than their tissue residence time. However, parameter estimation would then require accurate measurements of cell numbers within the tissue.

## Discussion

Our analysis indicates that CD4^+^ T_RM_ are not intrinsically long-lived, but instead are sustained by both self-renewal and supplementation from circulating precursors. By combining fate reporting methods with mathematical models, we also showed that it is possible to separately quantify the processes that underlie their persistence. We saw quite distinct contributions of recruitment and self-renewal of both subsets within skin and LP. The basis of this difference is unclear, but we speculate that the large antigenic burden within the small intestine drives the higher levels of T_RM_ recruitment and clonal erosion within the lamina propria. We showed that estimates of these quantities depend on the identity of any precursor, whose label kinetics propagate downstream into the population of interest; and the extent of any cell division that occurs around the time of differentiation or ingress. However, by using easily interpretable mathematical models, and assessing the statistical support for each, we were able to measure the support for different pathways and modes of recruitment into each subset.

The schematic in Fig. 1B illustrates a hypothetical example in which the frequency of YFP-expressing cells within a precursor declines. This trend is then reflected downstream in the target. However, in our experiments the kinetics of YFP in most of the putative precursors were quite flat (Fig. 2D). As noted above, these kinetics were likely due largely to the continued influx of new cells into circulating memory subsets, likely from naive precursors (Gossel *et al*., 2017, Hogan *et al*., 2019, Bullock *et al*., 2024). These had increasing levels of YFP (Fig. 2D, left panel) deriving from thymocytes that were dividing rapidly during treatment. YFP levels among naive T cells were also likely sustained to an extent by low-level residual labelling of thymic progenitors (Lukas *et al*., 2023). YFP expression was therefore not cleanly ‘washed out’ in the periphery. The data from the labelling of CD4-expressing cells were more informative for dissecting turnover; mTom^+^ cells were clearly diluted out of all peripheral populations by the descendants of mTom^−^ thymocytes.

In these reporter mice, YFP and mTom were induced quickly in all subsets to different degrees; therefore our inferences regarding precursor-target relationships weren’t informed by the initial levels of label in each. (For example, imagine a rapidly dividing target fed by a slowly dividing precursor; initially, YFP levels in the target would be higher than those in the precursor). The hierarchy of levels of label in different subsets *would* be informative if one expects targets to begin with no label at all; for instance, in the busulfan chimeric mouse system (Hogan *et al*., 2015) new, thymically derived ‘labelled’ (donor) cells progressively infiltrate replete ‘unlabelled’ (host) populations. In that case, one can immediately reject certain differentiation pathways by examining the sequence of accrual of donor cells in different subsets. In the systems we use here, information regarding lineage relationships is contained instead in the trends in YFP and mTom frequencies after treatment, because precursor kinetics must leave an imprint on the target (Fig. 1B). This information is particularly useful if two populations exhibit opposing trajectories – they are then unlikely to be immediately related.

On a technical note, in general one can reduce variation by comparing quantities derived from the same individual. We showed previously that in some situations, exploiting the within-mouse grouping of observations can reduce uncertainty and refine parameter estimates when modelling the dynamics of cell populations (Yates *et al*., 2007). We were not able to take this approach here, however, because the frequency of cells expressing YFP or mTom within a given subset in a particular mouse depends on the accumulated history of label in that subset’s precursor. We were unable to identify these trajectories for each animal, so we were obliged to use the population averages (Fig. 2D). To mitigate any biases this averaging might introduce, we fitted these empirical functions simultaneously with the models of label kinetics in the targets. This conservative strategy propagated the uncertainty in the precursor trajectories into our conclusions.

Our analysis was rooted in the observation that all CD4^+^ T cell subsets within these tissues were at steady-state. This dynamic equilibrium dictates that immigration of new cells must be accompanied by loss of existing ones. In the SPF mice studied here, these dynamic populations are likely specific for self or commensal antigens that are continuously expressed. It is possible that residence and interdivision times are distinct for T_RM_ that might not be replenished long-term from precursors, such as those generated in acute infections. Further, our simple model can explain the stable maintenance of T_RM_ numbers in healthy skin and LP without needing to invoke the concept of a homeostatic niche, such as a competitive limit to cell densities. In this model, any increase or decrease in the rate of influx into a tissue will simply lead to a new equilibrium at higher or lower cell densities, respectively. Indeed repeated vaccinia virus challenges can drive progressive increases in the numbers of virus-specific CD8^+^ T_RM_ in skin that are detectable for months (Jiang *et al*., 2012), and heterologous challenges appear not to erode pre-existing LCMV-specific CD8^+^ T_RM_ (Wijeyesinghe *et al*., 2021). However, whether the same flexibility manifests among CD4^+^ T_RM_ following repeated challenges is unclear.

Our goal here was to assess the support for external replenishment of effector-memory-like CD4^+^ T cells in bulk, and the dominant CD69^+^ T_RM_ subset. We found evidence that CD69^+^CD4^+^ T_RM_ in skin derive at least in part from a local CD69^−^ precursor. With richer phenotyping of tissue-localised cells, we could in principle use labelling trajectories to define more fine-grained differentiation pathways. One issue is that accurate measurement of label frequencies becomes more difficult as one resolves T_RM_ into smaller subsets; indeed we saw that label kinetics within the small CD69^−^ populations were relatively noisy. Another issue is that label kinetics operate on the timescales of the net loss rate of cell populations – death and onward differentiation, balanced by self-renewal. More frequent sampling would be required to resolve more transitory intermediates. Nevertheless, our study clearly exposes the highly dynamic nature of CD4^+^ T_RM_, sustained throughout life by both self renewal and continued influx from precursors. This tissue-specific influx, particularly if there are competitive limits to T_RM_ occupancy, may contribute to the differential longevity of immunity at different barrier sites.

## Acknowledgements

This work was supported by the National Institutes of Health (R01 AI093870 and U01 AI150680) and the Medical Research Council (MR/P011225/1).

## Methods

### Reporter mouse strains

Ki67^mCherry-CreERT^ Rosa26^RcagYFP^ (Ki67-DIVN) and CD4^CreERT^ Rosa26^RmTom^ (Cd4-FR) mice have been described previously (Bullock *et al*., 2024). Experimental Ki67-DIVN mice were homozygous for indicated mutations at both the Ki67 and Rosa26 loci. Experimental CD4-FR mice were heterozygous for the indicated mutations and both the CD4 and Rosa26 loci. Tamoxifen (Sigma) was diluted to 20mg/mL in corn oil (Fisher Scientific) and 100*μ*l (2mg) was administered to mice via oral feeding on day 0. Ki67-DIVN mice were injected with 2μg Thy1.2-BV510 (53-2.1) (BioLegend) 3 minutes prior to sacrifice to label T cells in the circulation. This protocol typically achieves >99% staining of circulating cells (Anderson *et al*., 2014) and less than 3% of cells recovered from our tissue samples were label positive (Fig. S1B), supporting the assumption of very low rates of false positive and false negative events. Mice were subsequently taken down at specified timepoints post tamoxifen treatment for organ collection.

### Cell preparation

All peripheral lymph nodes (LN), the small intestine (SI) and ear skin were taken from mice and processed into single cell suspensions. LNs were mashed through two pieces of fine gauze in a petri dish and washed with complete RPMI (ThermoFisher) supplemented with 5% FCS (ThermoFisher) (cRPMI). Cells were resuspended in cold PBS and counted using the CASY counter (Cambridge Bioscience). Peyer’s patches were excised from the antimesenteric side of the SI before being opened longitudinally and SI contents scraped out. SI pieces were placed in 20mL pre-warmed extraction media (cRPMI + 10mM HEPES (ThermoFisher) + 5mM EDTA (Sigma) + 1mM DTT (Abcam)) and incubated in 37^°^C shaking incubator for 30 minutes at 200rpm. Cells were filtered over 70*μ*m cell strainer (supernatant containing intra-epithelial lymphocytes not used), and SI pieces were placed in cold cRPMI supplemented with 10mM HEPES and allowed to settle. Supernatant was carefully poured off and SI pieces were finely minced, added to 20mL pre-warmed digestion media (RPMI + 10% FCS + 1.5mg/mL collagenase VIII (Sigma)) and incubated in 37^°^C shaking incubator for 30 minutes at 200rpm. After digestion, cells were passed through 70*μ*m cell strainer and washed with cRPMI + 10mM HEPES. The resulting cell suspension contains cells from the lamina propria (LP) of the SI. Ear skin was excised and separated into dorsal and ventral sides. Skin was finely minced, added to 4mL digestion buffer (cRPMI + 50mM HEPES + 37.5*μ*g/mL Liberase TL (Merck)+ 3.125mg/mL collagenase IV (ThermoFisher)+ 1mg/mL DNAse I (Merck)) and incubated in 37^°^C shaking incubator for 2 hours at 200rpm. Cells were filtered over 70*μ*m cell strainer and washed through with cRPMI.

### Flow cytometry

All cells isolated from skin and LP, and 5 × 10^6^ LN cells, were stained for analysis by flow cytometry. Cells were stained in 100*μ*l PBS with combinations of: CD8*α*-BUV395 (53-6.7), CD25-BUV395 (PC61), CD62L-BUV737 (MEL-14), TCR*γ*δ-BV421 (GL3), CD103-BV786 (M290) (all BD Biosciences); CD25-BV650 (PC61), CD103-BV421 (2E7), CD8*α*-BV570 (53-6.7), TCR*γ*δ-BV605 (GL3), NK1.1-BV650 (PK136), CD44-BV785 (IM7), CD45.1-BV605 (A20), CD4-BV711 (RM4-5), CD8b.2-APC (53-5.8) (all BioLegend); TCR*β*-PerCPCy5.5 (104) (Cambridge Bioscience); CD44-APCef780 (IM7) (eBioscience); and CD3-APCef780 (2C11), CD3-biotin (145-2C11), CD69-PeCy7 (H1.2F3), CD45.2-AF700 (104), nearIR live/dead, blue live/dead, yellow live/dead (all ThermoFisher). Cells were fixed for 20 minutes with IC fix (Invitrogen) and washed twice in FACS buffer (PBS + 0.1% BSA). Flow cytometric analysis was performed on either a Cytek Aurora spectral flow cytometer or a conventional BD LSR-Fortessa and analysed using FlowJo software (Treestar).

### Cell count calculations

Cell counts of LN populations were calculated by dividing the event count in a target population by the event count of live cells, multiplied by the total live cells in LN prep determined by CASY counter. Sizes of LP and skin populations were calculated using AccuCount (Spherotech) counting beads that were spiked into the sample prior to acquisition, as per manufacturer’s instructions.

### Mathematical modelling and Statistical Analysis

We fitted simultaneously the kinetically homogeneous mathematical model illustrated in Fig. 1B, and described in Supporting Information Text S1, to the time courses of frequencies of YFP^+^, Ki67^high^ in YFP^+^, Ki67^high^ in YFP^-^, and mTom^+^ cells in the target populations; and empirical descriptor functions describing the trajectories of the frequencies of YFP^+^ and mTom^+^ precursors (all data shown in Fig. 2C). We used a Bayesian estimation approach using Python and Stan (Stan Development Team, 2024) to perform these model fits. Code and data used to perform model fitting, details of the prior distributions for parameters, and figure generation notebooks are available at https://github.com/elisebullock/CD4TRM. Models were ranked based on information criteria estimated using the Leave-One-Out (LOO) cross validation method (Vehtari *et al*., 2017). See Supporting Information, Text S1 for full details.

