## Supporting Information for "Tissue resident memory CD4^+^ T cells are sustained by site-specific levels of self-renewal and continuous replacement"

### Supporting Figures

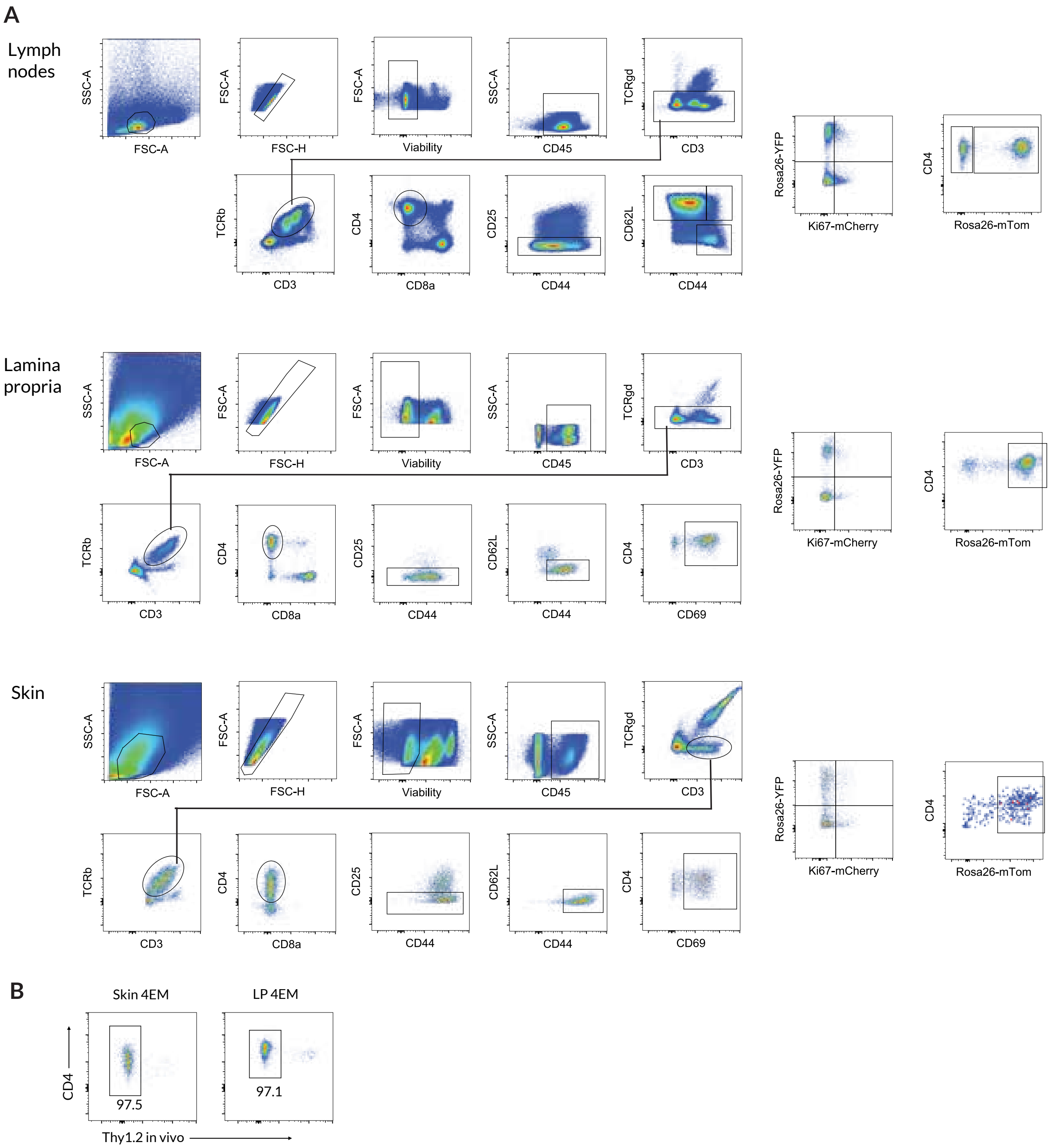


Figure S1: **A** Gating strategy for $CD4^+$ subsets in lymph nodes, LP and skin. **B** Tissue-localised $CD4^+$ EM subsets defined by protection from short-term *in vivo* labelling.

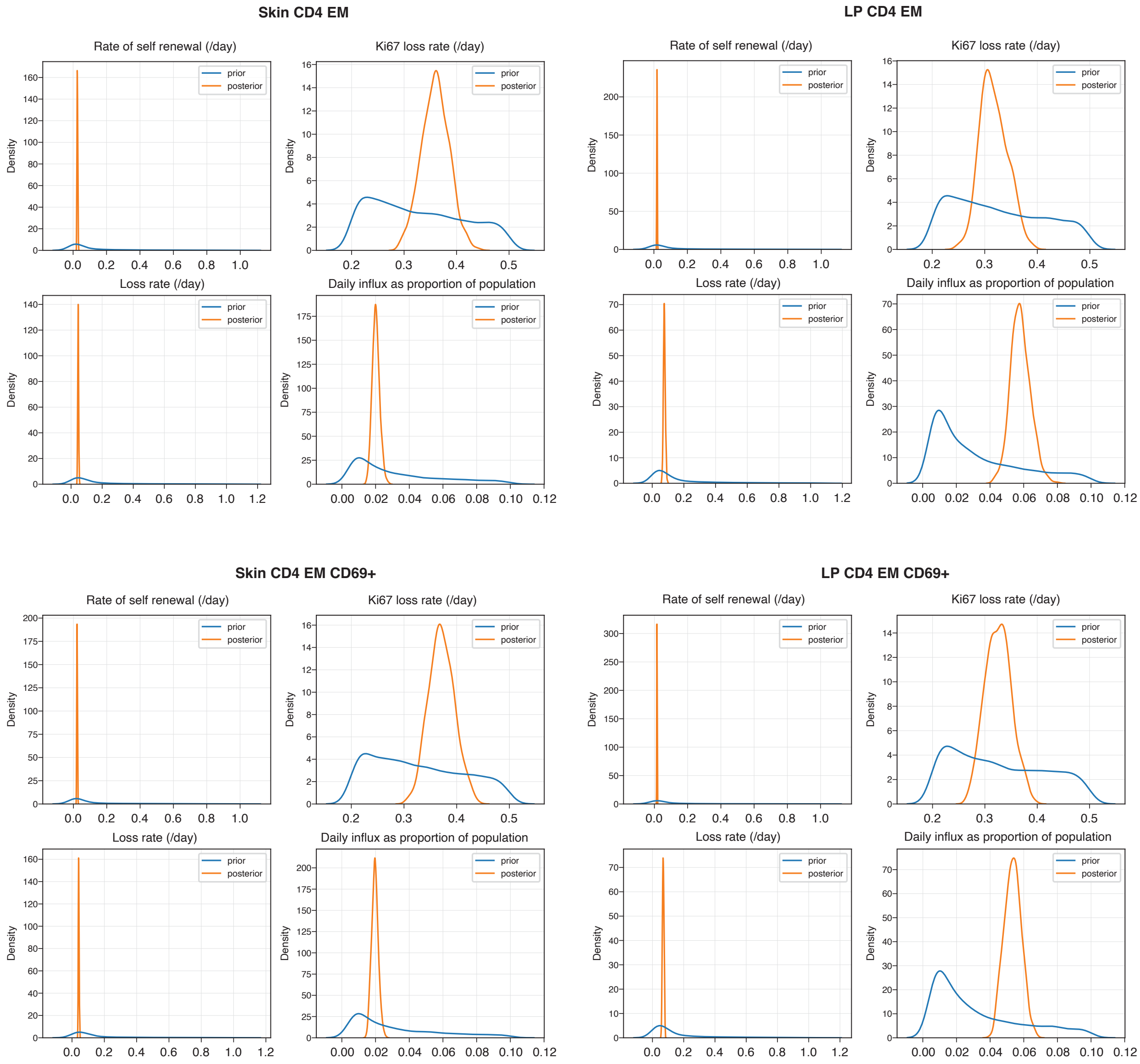


Figure S2: **Prior and posterior distributions of parameters for the best fitting models.** Prior distributions of all parameters were truncated lognormal distributions and here are shown as kernel-smoothed samples. Truncation limits: self-renewal rate $\alpha \in (0, 1)$; loss rate $\delta \in (\alpha, 5)$; rate of loss of Ki67 $\in (0.2, 0.5)$; daily influx as fraction of population size $\in (0.005, 0.1)$. The Ki67 loss rate was informed by our previous estimates (Bullock et al., 2024, Gossel et al., 2017). The constraint $\alpha < \delta$ ensures that the population is at a steady state. Upper bounds on division and loss rates translate to minimum interdivision and residence times of 24h and 4.8h respectively; the upper bound on the division rate was informed by Ki67 levels observed in tissues (see Text S2).

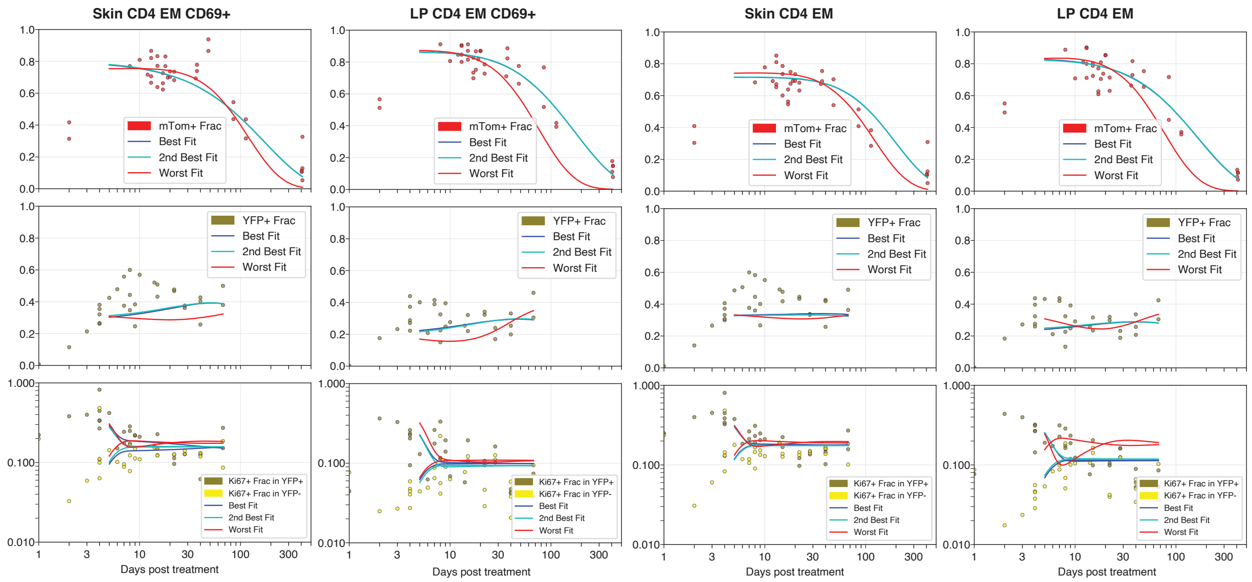


Figure S3: **Fits of top-, second-, and lowest-ranked models for each target population.** In cases where the best-fit (black) curves are not visible, they are closely overlaid by the blue curves (second-ranked model). Note that because each model was fitted to six time courses simultaneously, and fractional observations were logit-transformed to normalise residuals, apparently small visual differences in fits can nevertheless lead to substantial differences in model support.

### Supporting Tables

| Parameter | Tissue | Target | Recruitment mode | Precursor | Estimate |
|---|---|---|---|---|---|
| Mean residence time (days) | LP | $CD4^+$ EM | Quiescent ($Ki67^{low}$) | LN $CD4^+$ EM | 13 (11, 15) |
| | | $CD4^+CD69^+$ $T_{RM}$ | Neutral ($Ki67^{int}$) | LN $CD4^+$ EM | 14 (13, 16) |
| | Skin | $CD4^+$ EM | Division-linked ($Ki67^{hi}$) | LN $CD4^+$ EM | 22 (20, 24) |
| | | $CD4^+CD69^+$ $T_{RM}$ | Division-linked ($Ki67^{hi}$) | Skin $CD69^-$ | 24 (22, 27) |
| Mean interdivision time (days) | LP | $CD4^+$ EM | Quiescent ($Ki67^{low}$) | LN $CD4^+$ EM | 49 (44, 57) |
| | | $CD4^+CD69^+$ $T_{RM}$ | Neutral ($Ki67^{int}$) | LN $CD4^+$ EM | 66 (57, 74) |
| | Skin | $CD4^+$ EM | Division-linked ($Ki67^{hi}$) | LN $CD4^+$ EM | 39 (34, 46) |
| | | $CD4^+CD69^+$ $T_{RM}$ | Division-linked ($Ki67^{hi}$) | Skin $CD69^-$ | 46 (40, 53) |
| Clonal half-life (days) | LP | $CD4^+$ EM | Quiescent ($Ki67^{low}$) | LN $CD4^+$ EM | 12 (10, 14) |
| | | $CD4^+CD69^+$ $T_{RM}$ | Neutral ($Ki67^{int}$) | LN $CD4^+$ EM | 13 (11, 15) |
| | Skin | $CD4^+$ EM | Division-linked ($Ki67^{hi}$) | LN $CD4^+$ EM | 34 (29, 41) |
| | | $CD4^+CD69^+$ $T_{RM}$ | Division-linked ($Ki67^{hi}$) | Skin $CD69^-$ | 35 (30, 42) |
| Percent daily replacement by influx | LP | $CD4^+$ EM | Quiescent ($Ki67^{low}$) | LN $CD4^+$ EM | 5.7 (4.9, 6.9) |
| | | $CD4^+CD69^+$ $T_{RM}$ | Neutral ($Ki67^{int}$) | LN $CD4^+$ EM | 5.4 (4.5, 6.2) |
| | Skin | $CD4^+$ EM | Division-linked ($Ki67^{hi}$) | LN $CD4^+$ EM | 2.0 (1.7, 2.4) |
| | | $CD4^+CD69^+$ $T_{RM}$ | Division-linked ($Ki67^{hi}$) | Skin $CD69^-$ | 2.0 (1.7, 2.3) |
| Percent production from self-renewal | LP | $CD4^+$ EM | Quiescent ($Ki67^{low}$) | LN $CD4^+$ EM | 26 (22, 30) |
| | | $CD4^+CD69^+$ $T_{RM}$ | Neutral ($Ki67^{int}$) | LN $CD4^+$ EM | 22 (19, 27) |
| | Skin | $CD4^+$ EM | Division-linked ($Ki67^{hi}$) | LN $CD4^+$ EM | 55 (50, 62) |
| | | $CD4^+CD69^+$ $T_{RM}$ | Division-linked ($Ki67^{hi}$) | Skin $CD69^-$ | 52 (46, 59) |
| Ki67 Lifespan | LP | $CD4^+$ EM | Quiescent ($Ki67^{low}$) | LN $CD4^+$ EM | 3.4 (2.9, 4.0) |
| | | $CD4^+CD69^+$ $T_{RM}$ | Neutral ($Ki67^{int}$) | LN $CD4^+$ EM | 3.0 (2.7, 3.5) |
| | Skin | $CD4^+$ EM | Division-linked ($Ki67^{hi}$) | LN $CD4^+$ EM | 2.8 (2.5, 3.2) |
| | | $CD4^+CD69^+$ $T_{RM}$ | Division-linked ($Ki67^{hi}$) | Skin $CD69^-$ | 2.7 (2.4, 3.0) |

Table S1: MAP estimates and 95% credible intervals of parameters from the statistically favoured models. LP: Lamina propria. LN: Lymph node.

### Supporting Text

#### S1 Mathematical modelling

##### S1.1 The kinetics of mTom expression derived from Cd4-FR mice

We used a simple homogeneous ODE model to describe the kinetics of $\text{mTom}^+$ and $\text{mTom}^-$ cells by tracking their loss (per capita $\delta$), self-renewal (per capita rate $\rho$), and supplementation from a precursor population at constant total rate $\theta$ and with label content $f_{\text{mTom}}$, described empirically (see S1.3);

$$\frac{d}{dt}\text{mTom}^+ = \theta \cdot f_{\text{mTom}}(t) - (\delta - \rho) \cdot \text{mTom}^+$$
$$\frac{d}{dt}\text{mTom}^- = \theta \cdot (1 - f_{\text{mTom}}(t)) - (\delta - \rho) \cdot \text{mTom}^-$$

##### S1.2 Modelling the label trajectories derived from Ki67-DIVN mice

The model we used to describe data derived from the Ki67-DIVN mice was is similar to that above, with the same parameters $\theta$, $\delta$ and $\rho$, but included the kinetics of Ki67 expression (see equations below). Following mitosis, Ki67 protein levels continuously decrease. However, in standard flow cytometry analysis, cells are classified into two categories: $\text{Ki67}^{\text{high}}$ and $\text{Ki67}^{\text{low}}$. To model the transition between these states, we used the linear chain technique (MacDonald, 1978) and introduced intermediate Ki67 compartments as in previous studies (Bullock et al., 2024), allowing us to characterize the residence time within the $\text{Ki67}^{\text{high}}$ compartment as gamma-distributed with low variance, with 12 intermediate Ki67 levels ($\ell$). The rate at which $\text{Ki67}^{\text{high}}$ cells move between these intermediate stages is given by $\ell\beta$, such that the expected time that a cell spends within the $\text{Ki67}^{\text{high}}$ state before transitioning to the $\text{Ki67}^{\text{low}}$ state defined by the flow cytometric gating strategy is $1/\beta$. For each precursor/target pair, we examined three influx modes: one where new immigrant cells are $\text{Ki67}^{\text{low}}$ (quiescent), another where their Ki67 expression mirrors the precursor's, and a third where they are fully $\text{Ki67}^{\text{high}}$ after division. To account for this in the model, we used a parameter ($k_f$) which was set to either 0, the mean $\text{Ki67}^{\text{high}}$ fraction within the precursor, or 1, respectively. In the model, the subscripts denote the Ki67 state or expression level (0 is $\text{Ki67}^{\text{low}}$; $1 \ldots \ell$ are $\text{Ki67}^{\text{high}}$):

$$\frac{d}{dt}\text{YFP}^+_\ell = \theta \cdot f_{\text{YFP}}(t) \cdot k_f - (\delta + \ell\beta + \rho) \cdot \text{YFP}^+_\ell + 2\rho \sum_{j=0}^{\ell} \text{YFP}^+_j$$
$$\frac{d}{dt}\text{YFP}^-_\ell = \theta \cdot (1 - f_{\text{YFP}}(t)) \cdot k_f - (\delta + \ell\beta + \rho) \cdot \text{YFP}^+_\ell + 2\rho \sum_{j=0}^{\ell} \text{YFP}^-_j$$
$$\frac{d}{dt}\text{YFP}^i_j = \ell\beta \cdot \text{YFP}^i_{j+1} - (\delta + \rho + \ell\beta) \cdot \text{YFP}^i_j \qquad \text{for } i \in \{+,-\} \text{ and } 1 \leq j < \ell$$
$$\frac{d}{dt}\text{YFP}^+_0 = \theta \cdot f_{\text{YFP}}(t) \cdot (1 - k_f) + \ell\beta \cdot \text{YFP}^+_1 - (\delta + \rho) \cdot \text{YFP}^+_0$$
$$\frac{d}{dt}\text{YFP}^-_0 = \theta \cdot (1 - f_{\text{YFP}}(t)) \cdot (1 - k_f) + \ell\beta \cdot \text{YFP}^-_1 - (\delta + \rho) \cdot \text{YFP}^-_0$$

##### S1.3 Model Fitting

###### S1.3.1 Empirical functions describe label dynamics in precursor populations

We described the trajectories of YFP and mTom expression within the candidate precursors with simple functions based on exponentials. In most populations, the labelled fractions ($f_{\text{mTom}}$ and $f_{\text{YFP}}$), both decreased over time, each described with a curve of the functional form

$$f(t) = ae^{-bt}. \tag{1}$$

We used a saturating exponential increase function for the YFP$^+$ fraction within LN-derived naive CD4 T cells,

$$f(t) = a(1 - e^{-bt}). \tag{2}$$

For each precursor we fitted these functions to the data alone using FME (Soetaert and Petzoldt, 2010) and in Python using the curvefit() function from the package SciPy (Virtanen et al., 2020). These fits are shown in Fig. 2D. The best-fit parameters (see table below) and associated uncertainties were used to inform the priors for fitting precursor and target trajectories simultaneously using our Bayesian approach.

| Population | $a_y$ | $b_y$ | $a_t$ | $b_t$ | $k_f$ |
|---|---|---|---|---|---|
| LN CD4 CM | 0.286 | 0.001 | 0.857 | 0.011 | 0.2 |
| LN CD4 EM | 0.289 | 0.002 | 0.833 | 0.007 | 0.12 |
| LN CD4 Naive | 0.372 | 0.081 | 0.919 | 0.030 | 0.16 |
| SK CD69$^-$ | 0.258 | 0.001 | 0.542 | 0.007 | 0.21 |
| LP CD69$^-$ | 0.449 | 0.007 | 0.457 | 0.007 | 0.0047 |

Fitted estimates of parameters defining the empirical descriptions of precursor label trajectories ($y$ for YFP, and $t$ for mTom). The $k_f$ term describes the mean Ki67$^{\text{high}}$ fraction within the precursor population, as described in S1.2.

##### S1.3.2 Initialising the system before describing post-treatment kinetics of labelled populations

Motivated by observations (Fig. 2A in the text) we assumed all target populations in skin and LP were at homeostatic equilibrium, and with constant levels of Ki67. To ensure the system was at this equilibrium before tamoxifen treatment, for each set of parameter values we used the steady state assumption and observed Ki67 fraction (estimated by taking the mean of the observed values in each population) to eliminate two parameters from the model (see below). We then utilized the Runge-Kutta 45 ODE solver within Stan, setting all species to zero, and running for $10^4$ days to drive the system to this prescribed steady state. The labeled fractions of cells in the target population at d5 post treatment were free parameters $\epsilon_{\text{YFP}}$ and $\epsilon_{\text{mTom}}$, which reflected the cumulative effect of label induction up to that point. Cells were then distributed into labelled and unlabelled populations using these parameters.

##### S1.3.3 Modeling noise and mouse-to-mouse variation

In addition to the empirically described trajectories of label content within each precursor population, from the model we derived the four proportions of interest for the target: the mTom$^+$ fraction, the YFP$^+$ fraction, and the Ki67$^{\text{high}}$ fractions within YFP$^{+/-}$ cells. As all of these observations lay within the interval $(0, 1)$, we logit-transformed them to normalise residuals. The normally distributed noise within each dataset was defined with six separate standard deviations, $\sigma$, each themselves described with normal priors. Fits were obtained using all six timeseries simultaneously (summing the log likelihoods). For performing statistical comparisons of different precursor-target relationships using LOO-IC, we used only the log-likelihoods for the four quantities derived from the target cell kinetics; this ensured we were comparing descriptions of the same observations.

##### S1.3.4 Prior information on parameters used for Bayesian distributions

By assuming that the population was at steady state and that the Ki67 fraction was constant, we could eliminate two of the five parameters ($\theta$, $\beta$, $\rho$, $\delta$ and $k_F$, the fraction of immigrant cells that are assumed to enter as Ki67$^{\text{high}}$).

All parameters were sampled from a range of values such that populations remained positive and finite. The prior on the Ki67 lifetime $1/\beta$ peaked at 3.25 days and constrained to remain between 2.5-4.5 days, based on previous studies. Based on previous studies (Gossel *et al.* eLife 2017; Bullock *et al.* Plos Biology 2024) the influx rate $\theta$ was constrained to be between 0.05% and 10% of the total pool size. We set broad priors on the initialization induction efficiencies of YFP and mTom ($\epsilon_{\text{YFP}}$ and $\epsilon_{\text{mTom}}$), with upper limits of 1.

All code and data can be obtained at https://github.com/elisebullock/CD4TRM.

#### S2 Estimating the average cell residence times using Ki67

Consider a cell population at steady state numbers $N$ with cells dividing at an average *per capita* rate $\rho$, leaving the population through death, egress, or differentiation at average *per capita* rate $\delta$, and being replenished from a precursor population at total rate $\theta$;

$$\frac{dN}{dt} = \theta + (\rho - \delta)N = 0 \implies \rho N + \theta = \delta N. \tag{3}$$

Assume a fraction $f$ of immigrants enter the population as Ki67$^{\text{high}}$. We know that Ki67 is expressed for a time $T \simeq 3-4$ days during and after division. Then the number of cells within the population as a whole that are Ki67$^{\text{high}}$, $K^+$, is constant and equal to the number that were produced by division or entered as Ki67$^{\text{high}}$ in the last $T$ days, and did not divide and survived to the present;

$$K^+ = \int_0^T (2\rho N + f\theta)\, e^{-(\delta+\rho)(T-s)} ds = \frac{2\rho N + f\theta}{\delta + \rho}(1 - e^{-(\delta+\rho)T}), \tag{4}$$

The Ki67$^{\text{high}}$ proportion observed within the total population at any time is therefore also constant and equal to $k = K^+/N$.

$$k = \frac{2\rho + f\frac{\theta}{N}}{\delta + \rho}(1 - e^{-(\delta+\rho)T}). \tag{5}$$

In the general case where influx is substantial, we need an independent estimate of the daily proportional replacement through influx, $\theta/N$, which by eqn. 3 is equal to the daily net loss rate $\lambda = \delta - \rho$. Then, eliminating the division rate $\rho$,

$$k = \frac{2(\delta - \lambda) + f\lambda}{2\delta - \lambda}(1 - e^{-(2\delta-\lambda)T}). \tag{6}$$

Given an observed value of $k$, the known value of $T$, an estimate of the replacement (influx) rate $\lambda$, and a value for $f$ (reflecting the extent to which incoming cells are recently divided) eqn. 6 can be solved numerically for the cell lifespan $\tau = 1/\delta$.

We can consider some simpler limiting cases. If most cell production occurs through self renewal rather than influx, then $\rho \simeq \delta$ and $\lambda \simeq 0$. Eqn. 6 then becomes $k \simeq (1 - e^{-2T/\tau})$. This yields a simple formula that relates the Ki67 fraction within a closed or nearly-closed population to the mean lifespan of its constituent cells;

$$\tau = \frac{-2T}{\ln(1-k)} \simeq \frac{2T}{k} \text{ when } k \text{ is small.} \tag{7}$$

If influx cannot be disregarded, but we have an indication that cell lifespans are much longer than the duration of Ki67 expression, $\tau \gg T$ (which will be likely if $k$ is small, suggesting slow turnover), then $e^{-(2\delta-\lambda)T} \simeq 1 - T(2\delta - \lambda)$ and eqn. 6 gives

$$k \simeq T(\lambda(f-2) + 2/\tau) \implies \tau \simeq \frac{2T}{k + \lambda T(2-f)}. \tag{8}$$

Rough bounds on the mean cell lifespan are then

$$\frac{2T}{k + 2\lambda T} < \tau < \frac{2T}{k + \lambda T}, \tag{9}$$

spanning the cases $f = 0$ (all newly recruited cells are quiescent) and $f = 1$ (all new immigrants are recently divided; for example, having recently been antigen stimulated). A value of $f$ between 0 and 1 might also arise if the mean duration of Ki67 expression on new immigrants is shorter than the 3-4 days expected on a self-renewing cell; reflecting the possibility that they last divided as precursors less than $T$ days before entering the population.
